# Heat Shock Protein 27 Immune Complex Altered Signaling and Transport (ICAST): Novel Mechanisms of Attenuating Inflammation

**DOI:** 10.1101/2020.05.31.126581

**Authors:** Chunhua Shi, Jingti Deng, Michael Chiu, Yong-Xiang Chen, Edward R. O’Brien

## Abstract

Blood levels of heat shock protein (HSP27) and natural IgG auto-antibodies to HSP27 (AAbs) are higher in healthy controls compared to cardiovascular disease patients. Vaccination of mice with recombinant HSP25 (rHSP25, murine ortholog of human rHSP27) increased AAb levels, attenuated atherogenesis and reduced plaque inflammation and cholesterol content. We sought to determine if the HSP27 immune complex (IC) altered MΦ inflammation signaling (Toll Like Receptor 4; TLR4), and scavenger receptors involved in cholesterol uptake (SR-AI, CD-36). Combining a validated polyclonal IgG anti-HSP27 antibody (PAb) with rHSP27 enhanced binding to THP-1 MΦ cell membranes and activation of NF-κB signaling via TLR4, competing away LPS and effecting an anti-inflammatory cytokine profile. Similarly, adding the PAb with rHSP27 enhanced binding to SR-AI and CD-36, as well as lowered oxLDL binding in HEK293 cells separately transfected with SR-AI and CD-36, or THP-1 MΦ. Finally, the PAb enhanced the uptake and internalization of rHSP27 in THP-1 MΦ. Thus, the HSP27 IC potentiates HSP27 cell membrane signaling with receptors involved in modulating inflammation and cholesterol uptake, as well as HSP27 internalization. Going forward, we will explore HSP27 Immune Complex Altered Signaling and Transport (ICAST) as a new anti-inflammatory therapeutic strategy *in vivo*.

## Introduction

Atherosclerosis is a chronic inflammatory disease of the artery wall that is characterized by the accumulation of lipids in macrophages (MΦ) (1). Heat Shock Protein 27 (HSP27) is a member of the small heat-shock protein family that is primarily known as an intracellular chaperone, and more recently for its extracellular, anti-inflammatory roles (2). Our laboratory demonstrated that HSP27 is a potential biomarker for atherosclerosis, with its expression attenuated in the presence of atherosclerosis (3) – an observation that 4 other groups confirmed using objective (proteomic discovery) approaches (4-7). In humans, low serum HSP27 levels are associated with the presence of coronary artery disease and a higher 5-year likelihood of future adverse clinical events (e.g., heart attack, stroke, cardiovascular death) (8). Cross-breeding mice that over-express HSP27 with atherosclerosis-prone *ApoE*^*-/-*^ mice we demonstrated that HSP27 reduces serum and plaque cholesterol levels, vessel wall inflammation and unstable lesion formation (8-10). *In vitro*, recombinant HSP27 reduced MΦ uptake of modified low density lipoprotein (LDL), inhibited secretion of the pro-inflammatory cytokine interleukin-1β (IL-1β) and promoted the release of anti-inflammatory IL-10 (9). Moreover, HSP27 binds to scavenger receptor AI (SR-AI) thereby reducing lipid uptake and the conversion of MΦ into foam cells (9).

Recently, we discovered that blood levels of natural IgG (but not IgM) auto-antibodies to HSP27 (AAbs) are higher in healthy control subjects compared to cardiovascular disease patients (11). Vaccinating *ApoE*^*-/-*^ mice with recombinant HSP25 (rHSP25; murine ortholog of human rHSP27) increased AAb levels, attenuated atherogenesis, reduced plasma cholesterol levels as well as the abundance of plaque inflammation and cholesterol content. As the role of these AAbs in athero-protection is not yet clear, we began our studies by trying to determine how the immune complex (IC) that forms between HSP27 and AAbs alters: i) MΦ signaling, looking specifically at the NF-κB inflammation pathway, and ii) the uptake of oxidized LDL (oxLDL). We used a validated rabbit anti-HSP27 polyclonal antibody (PAb) that shares the same epitope mapping pattern as human plasma derived AAbs (11). Herein, we show that compared to HSP27 alone, the HSP27 IC shows enhanced cell membrane interactions that result in altered signaling with key receptors involved in inflammation (Toll Like Receptor 4; TLR4) and oxLDL uptake via the scavenger receptors, SR-AI and CD-36.

## Methods and Materials

### Cell Culture Experiments

a. The THP-1 monocytic MΦ cell line was obtained from American Type Culture Collection (ATCC# TIB-202; Manassas, VA) and incubated at 37°C and 5% CO_2_ in complete RPMI 1640 growth medium supplemented with 10 U/ml penicillin and streptomycin, 1mM sodium pyruvate (11965092; Thermo Fisher Scientific, Burlington, ON) and heat inactivated 10% Fetal Bovine Serum (FBS; Gibco, Burlington, ON). These undifferentiated mononuclear cells (between passage 3 and 16) were seeded in 24-well plates at concentration of 1×10^6^ cells / well, and stimulated with 50 ng/ml phorbol 12-myristate 13-acetate (PMA; Sigma-Aldrich, Oakville, ON) to induce differentiation to MΦ-like cells (12).
b. HEK-Blue™ Null1-v and HEK-Blue™-TLR4 (stably expressing TLR4, MD-2 and CD14 co-receptor genes) cell lines were obtained from Invivogen (SanDiego, CA, USA). HEK-Blue™ SR-AI and HEK-Blue™ CD36 were prepared by stably transfecting the parental cell line HEK-Blue™ Null1-v with plasmid pCMV/Hygro-SR-AI or pCMV/Hygro-CD36 (Sino Biological Inc., North Wales, PA) and selecting using the addition of 100 μg/mL hygromycin. The medium for all the HEK-Blue™ cell lines was DMEM, supplemented with 4.5 g/l glucose, 2-4 mM L-glutamine with heat-inactivated 10% (v/v) FBS, 50 U/ml penicillin, and 50 μg/ml streptomycin. All HEK-Blue™ cell lines contain an inducible secreted embryonic alkaline phosphatase (SEAP) reporter gene under the control of an IFN-β minimal promoter fused to five NF-κB and AP-1-binding sites (Invivogen).
c. THP1 XBlue™ cells (Invivogen) that were maintained and subcultured in complete RPMI-1640 media supplemented with Zeocin (200 μg/mL) as a selective antibiotic.

### Recombinant Protein preparation

Plasmids encoding HIS-tagged full-length recombinant HSP27 (rHSP27) or the C-terminal, biologically inactive (rC1) fragment of HSP27 (AA93-205) (**Supplemental Fig. 1A**) were constructed using a pET-21a vector, and the plasmids were transformed into an *Escherichia coli* expression strain (DE3) as described previously (13). Recombinant proteins were purified with a Ni-NTA resin and Q-Sepharose™ (GE Healthcare, Mississauga, ON). Endotoxin was removed using the Pierce™ High-Capacity Endotoxin Removal Resin (ThermoFisher Scientific; Burlington, ON). The purity of the final recombinant proteins was more than 99% with an endotoxin concentration lower than 2 units/mg protein by Limulus Amebocyte Lysate PYROGENT™ 125 Plus (Lonza; Walkersville, MD).

### Generation and Validation of Anti-HSP27 IgG polyclonal antibody

A rabbit polyclonal IgG antibody (PAb) mimicking the human HSP27 autoantibody was produced according to the standard procedure by Cedarlane Laboratories (Burlington, ON) and the standard requirements of the Canadian Council on Animal Care. The production and validation of this PAb is previously described (11). To verify the fidelity of this PAb we compared its immunoreactivity with a commercial (goat) anti-HSP27 antibody using Western blotting of rHSP27 and rHSP25. As well, we used a spot blot epitope mapping technique to compare the immunoreactivity of the PAb *vs*. naturally occurring anti-HSP27 antibodies found in human plasma. Specific 15-mers of the linearized protein reacted with the PAb in a pattern that was identical to that of the naturally occurring AAbs – whether they were derived from the blood of healthy control or cardiovascular disease subjects. The binding of the PAb to these epitopes was abrogated by the addition of rHSP27 (100 μg/ml *vs*. an irrelevant anti-human IgG negative control), thereby indicating that epitopes in the full-length rHSP27 were similar to those on the spot blot.

### Fast Protein Liquid Chromatography (FPLC)

The molecular size of the rHSP27 and rC1 proteins with or without the PAb (using ratios of 1:1 and 1:5 for Hsp27:PAb; wt:wt) was determined by size exclusion chromatography in AKTA Primer Plus fast protein liquid chromatography system (GE Healthcare) with Superose™ 6 10/30 GL Column (GE Healthcare; **Supplemental Fig. 1B**). Samples were diluted in phosphate buffered saline (PBS) to 20 μg/ml and the PAb was mixed with rHSP27 for 30 mins. A final sample volume of 0.2 ml was loaded on the columns, the fractions were eluted with 0.2 ml/min PBS buffer and absorption was monitored at 280 nm. A standard curve was constructed using a mix of blue dextran (2,000 kD, Millipore Sigma, Oakville, ON), apoferritin (443 kD, Millipore Sigma,), Alcohol Dehydrogenase from yeast (150 kD; Millipore Sigma) and bovine serum albumin (BSA; 66kD), as well as human LDL and HDL (Biomedical Technologies).

### Physical Interaction Between HSP27 and PAb

Isothermal Titration Calorimetry (ITC) was used to gauge the strength of the interaction between HSP27 and the PAb, as reflected by the energy released from the interaction of two components (14-17). ITC experiments were performed using an ITC200 micro-calorimeter (MicroCal, North Hampton MA USA) with a reaction cell and reference cell volume of 200 μL and a syringe volume of 40 μL. PAb and rHSP27 were prepared in PBS (pH 7.4) with dialysing membranes utilized to remove impurities. All solutions were degassed for 20 mins at 37°C prior to loading into the cells or syringe. The ligand was titrated into the reaction cell in 2 μL aliquots for a duration of 4 secs with 160 secs intervals in between additions. A total of 20 injections per experiment were used, with the first injection volume of 0.2 μL for 0.4 secs to remove any extraneous bubbles from the syringe. In order to account for the heat of dilution, PAb was also titrated into PBS and subtracted from the data set. MicroCal Origin 7 software (https://www.originlab.com) was utilized to perform data analysis and fitment of titration curves to a one binding site model (**Supplemental Fig. 1C**).

### Membrane Binding Assay

To determine the affinity of HSP27 to bind the cell membrane with or without the PAb, THP-1 MΦ were incubated with biotin-labeled rHSP27 or rC1 plus streptavidin-horse radish peroxidase (strep-HRP) at 4°C for 30 mins to prevent endocytosis of the reagents. Cells were then washed thrice with cold PBS. The substrate 3,3′,5,5′-tetramethylbenzidine (TMB; Millipore Sigma) was then added to detect the retained HRP activity for 10 mins. The reaction was stopped by 2N H_2_SO_4_ and the color was quantified at 450 nm using a Synergy Mx plate reader (BioTek; Winooski, VT).

### [rHSP27 + PAb] Binds TLR4

Previously, we showed that HSP27 (alone) binds TLR4, and activates the NF-κB pathway in a TLR4 dependent manner (18). We now sought to determine if addition of the PAb enhances the binding of rHSP27 to TLR4 (see schematic overview: **Supplemental Fig. 2A**). First, we set up a system to pull down the TLR4 protein. An anti-TLR4 capture antibody (100 ng/well in carbonate-bicarbonate buffer; R&D Systems, Minneapolis, MN) with demonstrable specificity for TLR *in vitro* (see Western blot: **Supplemental Fig. 2B**) was coated onto NUNC maxisorp plates covered with an adhesive plastic and incubated overnight at 4°C. The coating solution was then removed, and the plate was blocked by adding 200 μl 1% BSA in phosphate buffered saline tween (PBST) per well for at least 2 hrs at room temperature (RT). Next, we harvested TLR4 from THP-1 MΦ as follows. Approximately 5×10^7^ THP-1 MΦ were re-suspended in 1 ml of 2% Trition-100/PBST, vigorously shaken for at least 2 hrs at 4°C, followed by the addition of 4 ml 1% BSA/PBS before being homogenized by sonication for 10 secs. After washing twice with 200 μl PBST, 100 μl of THP-1 MΦ lysate was added to the plate for an additional 1 hr incubation at RT to pull down TLR4. Finally, 100 μl of PBST containing 1% BSA with 1 μg/ml biotinylated rC1 or rHSP27 in the presence or absence of 5 μg/ml PAb and 1 μg/ml strep-HRP was then applied for the interaction assay. After a 1 hr incubation at RT, the plate was washed 3 times with 200 μl/well, mixed with 100 μl of TMB and incubated for 10 mins at RT for development of the blue color. The reaction was stopped by the addition of 50 μl 2M H_2_SO_4_ and the optical absorbance at 450 nm was read using a Synergy Mx plate reader (**Fig. 1B**).

**Fig. 1.**
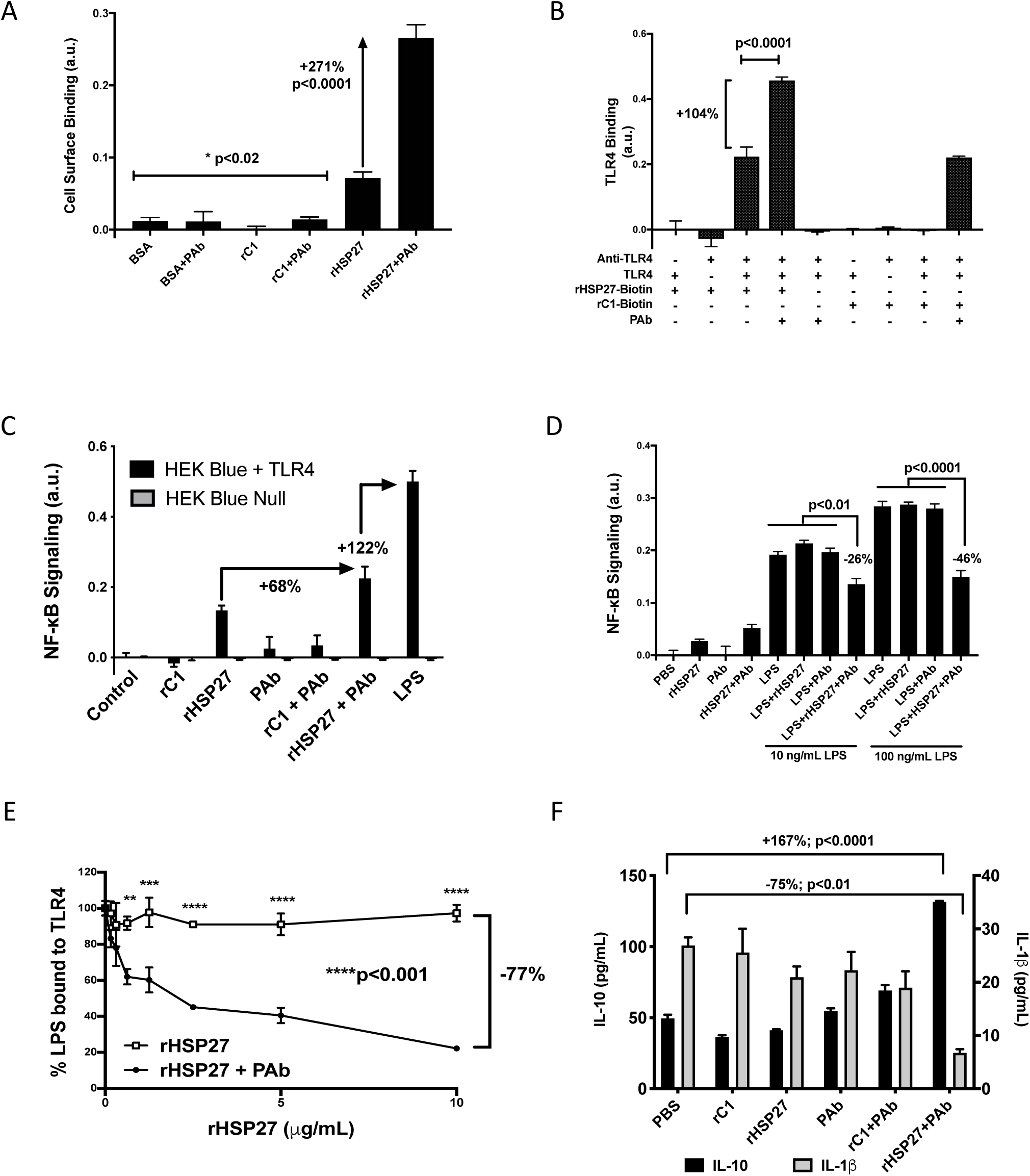
Hsp27 autoantibodies potentiate the anti-inflammatory effect of Hsp27. A) Cell surface binding assay using THP-1 MΦ, showing that PAbs promotes a 271% increase in rHSP27 binding to the cell surface (p<0.0001). B) *In vitro* binding assay using THP-1 MΦ cell lysates, showing that the presence of PAB strengthens rHSP27 binding to TLR4 by 104% (p<0.0001). C) NF-kB SEAP reporter gene assay of THP1-XBlue™ MΦ monocytes treated for 24 hr with different combinations of rHSP27 (1 μg/ml), rC1 (1 μg/ml), and PAb (5 μg/ml). The addition of the PAb to rHSP27 increased NF-kB signaling by 68% compared to HSP27 alone, but paled in comparison to LPS. Cells devoid of TLR4 showed only background NF-κB signals in response to all treatments. D) NF-κB reporter assay using THP1 XBlue™ MΦ shows that [rHSP27 + PAb] increased NF-κB signaling compared to rHSP27 alone. More importantly, the HSP27 IC attenuated LPS-mediated activation of NF-κB by 26% and 46% for LPS concentrations of 10 and 100 ng/ml (p<0.01 and <0.0001; respectively), thereby highlighting a competition between LPS and the HSP27 IC for TLR4. E) The HSP27 IC competitively inhibits the binding of biotinylated LPS to TLR4 by as much as 77% compared to HSP27 alone (p<0.001). F) In the culture media of THP-1 MΦ treated with LPS, cotreatment with [rHSP27 + PAb] decreased the concentration of IL-1β by 75% (p<0.01; left axis) and increased IL-10 by 167% (p<0.0001; right axis).

### [rHSP27 + PAb] Activates NF-κB via TLR4

To test the impact of the HSP27 IC on inflammatory signaling we conducted two experiments, both involving cells containing an NF-κB reporter construct. These experiments were done with the previously reported knowledge that the amount of endotoxin contamination in these recombinant proteins is measurably low, and of no functional consequences – as the addition of polymyxin B did not alter the NF-κB signal (12, 13). Lipopolysaccharide (LPS; from E. coli O111:B4, Cedarlane;) the principal component of Gram-negative bacteria that activates the innate immune system via TLR4, was used as a comparative positive control for both experiments.

The first experiment (**Fig. 1C**) sought to confirm the specificity and requirement for signaling of TLR4 in NF-κB signaling; hence, we used Human Embryonic Kidney (HEK) Blue™ Null1-v (i.e., devoid of TLR4) and HEK Blue™ TLR4 (stably expressing TLR4, MD-2 and CD14 co-receptor genes) cell lines (Invivogen). Cells were subject to treatment for 24 hrs with rHSP27 (1 μg/mL), rC1 (1 μg/mL), PAb (5 μg/mL), rHSP27 (1 μg/mL) or rC1 (1 μg/mL) plus PAb (5 μg/mL) or LPS (10 ng/mL). The conditioned media from each treatment group was analyzed for the presence of SEAP using QUANTIBlue™ detection reagent (Invivogen) according to the manufacturer’s instructions. Briefly, QUANTI-Blue™ detection reagent was mixed with cell supernatant (10:1) and incubated at 37°C for up to 1hr. Optical absorbance at 620 nm was then measured using the BioTek Synergy Mx microplate reader. Media only controls (no cells) were assayed to ensure that there was no endogenous SEAP activity in the treatment media. For the second experiment (**Fig. 1D**), we tested NF-κB activation in THP1 XBlue™ cells (Invivogen) treated with PBS as a control, and various combinations of rHSP27, PAb, and two concentrations of LPS (10 and 100 ng/ml), before assaying the conditioned media for SEAP.

### LPS Competitive Binding Assay

To compare the avidity of HSP27 with or without the PAb in competing with TLR4 we established an LPS competitive binding assay. After pulling down TLR4, 10 ng/ml of biotinylated LPS (LPS-EB from E. coli O111:B4; Cedarlane) was mixed with rHSP27 (concentration varying from 0 to 10 μg/ml) in the presence or absence of PAb (5 μg/ml) plus strep-HRP (1ug/ml). The binding of biotinylated LPS to TLR4 was then quantified using a Synergy Mx plate reader (BioTek) to record optical absorbance at 450 nm.

### Cytokines assays

For 2 hrs, THP-1 MΦ were treated with LPS (10 ng/ml) plus rHSP27 or rC1 (1 μg/ml) with or without PAb (5μg/ml). PBS or PAb alone constituted additional control treatments. The culture media was then harvested and the concentrations of human IL-1β and IL10 were measured using ELISAs (DY201 and DY217B, R&D Systems, Oakville, ON).

### [rHSP27 + PAb] Binds Scavenger Receptors

Previously, we showed that HSP27 (alone) binds SR-AI (9) and we now sought to determine if the HSP27 IC also binds SR-AI as well as the closely related scavenger receptor CD-36. SR-AI and CD-36 proteins were pulled down from MΦ membrane extracts as described above for TLR-4 using anti-SR-A (R&D System, 1:1000) and anti-CD36 (R&D System, 1:1000) antibodies with demonstrable affinities (see Western blots: **Supplemental Fig. 2B**). Next, PBST (100 μl) containing 1% BSA, 1 μg/ml biotinylated rC1 or rHSP27 in the presence or absence of 5 μg/ml PAb and 1 μg/ml strep-HRP were added to determine if there was an interaction between the HSP27 IC and the pulled-down scavenger receptors, as reflected by quantification of HRP activity as described for the TLR4 binding assay (**Fig. 2A, 2B**).

**Fig. 2.**
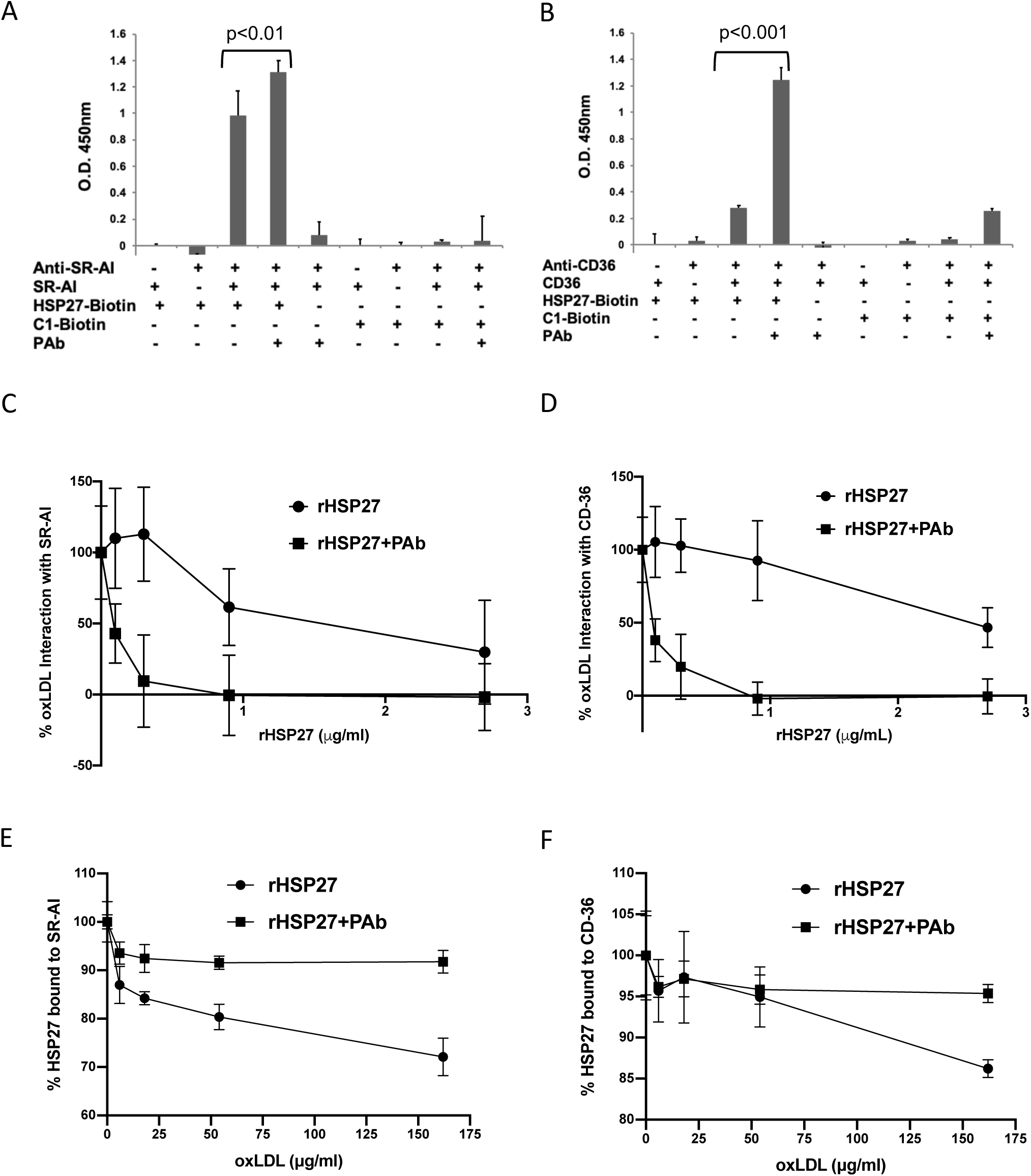
HSP27 IC competes with oxLDL to bind scavenger receptors SR-AI and CD36. A-B) *In vitro* binding assay using THP-1 MΦ cell lysates to show that the presence of PAb strengthened the binding of rHSP27 to either SR-AI or CD36. C-D) The interaction between oxLDL and SR-AI or CD36 is efficiently blocked by the HSP27 IC, but not HSP27 alone. E-F) Unlike the HSP27 (alone), the interaction between rHSP27 IC and either SR-AI or CD36 is effectively unchanged despite increasing the concentration of oxLDL.

### Competitive Binding Assays for Scavenger Receptors

Competitive inhibition assays were devised to compare the binding of oxLDL to either SR-AI or CD-36 in the presence of either rHSP27 or [rHSP27 + PAb]. As described for the TLR4 competitive binding assay described above, SR-AI and CD36 were separately pulled down from MΦ cell lysates, and the binding of biotinylated oxLDL to these receptors was measured while the concentration of rHSP27 was varied (0 to 2.7 μg/ml) in the presence or absence of PAb (**Fig. 2C, 2D**). The opposite competition assay (**Fig. 2E, 2F)** was also devised, looking at the binding of biotinylated rHSP27 (with or without PAb) to either SR-AI or CD-36 while the concentration of oxLDL was varied (0 – 164 μg/ml). Strep-HRP (1ug/ml) was then applied to create a quantifiable colorimetric reaction product reflective of the degree of interaction with either scavenger receptor.

### MΦ Flow Cytometry Assay for DiI-oxLDL Uptake

Uptake of oxLDL was analyzed using the following cells and assays:

i. To test the hypothesis that PAb also potentiates HSP27-mediated oxLDL uptake via scavenger receptors, HEK-Blue SR-AI and HEK-Blue CD36 cells were prepared from the parent cell line (HEK-Blue™ Null1-v) by stable transfection plasmid pCMV/Hygro-SR-AI or pCMV/Hygro-CD36 (Sino Biological Inc.) using Hygromycin as the selective antibiotic. The HEK-Blue cell lines were grown in DMEM, 4.5 g/l glucose, 2-4 mM L-glutamine with 10% FBS (heat activated), penicillin, streptomycin and Normocin™ (Invivogen). To confirm the expression of the transiently transfected scavenger receptors Western blotting and FACS were used, with the results summarized in **Supplemental Fig. 3A – 3D**. All cells were grown to 80% confluence, then treated with rHSP27 or rC1 with or without PAb (or PAb alone as a control) and incubated with 1µg/ml Dil-oxLDL for 2h. Cells were washed in PBS twice, detached and subsequently washed in 1× PBS and analyzed by flow cytometry with the phycoerythrin (PE) channel on 532 nm using a BD LSRII flow cytometer (BD Biosciences, San Jose, CA). The results are summarized (**Fig. 3A**) using data from one of three representative experiments, each quantifying: a) the percentage of cells that were fluorescent, and b) the mean fluorescence intensity (MFI) which represents the amount of DiI-oxLDL uptake for a particular experimental condition. **Fig. 3.**
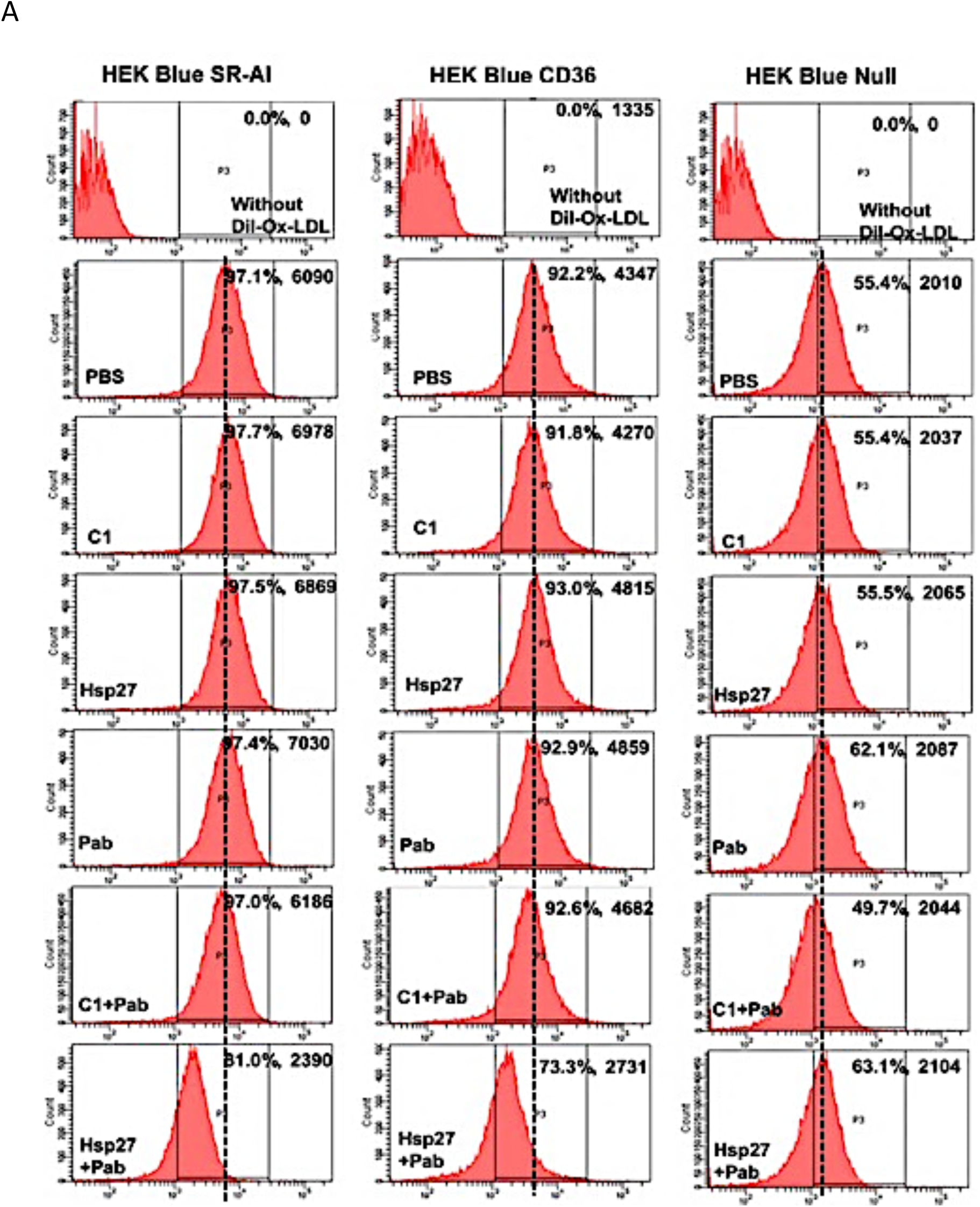
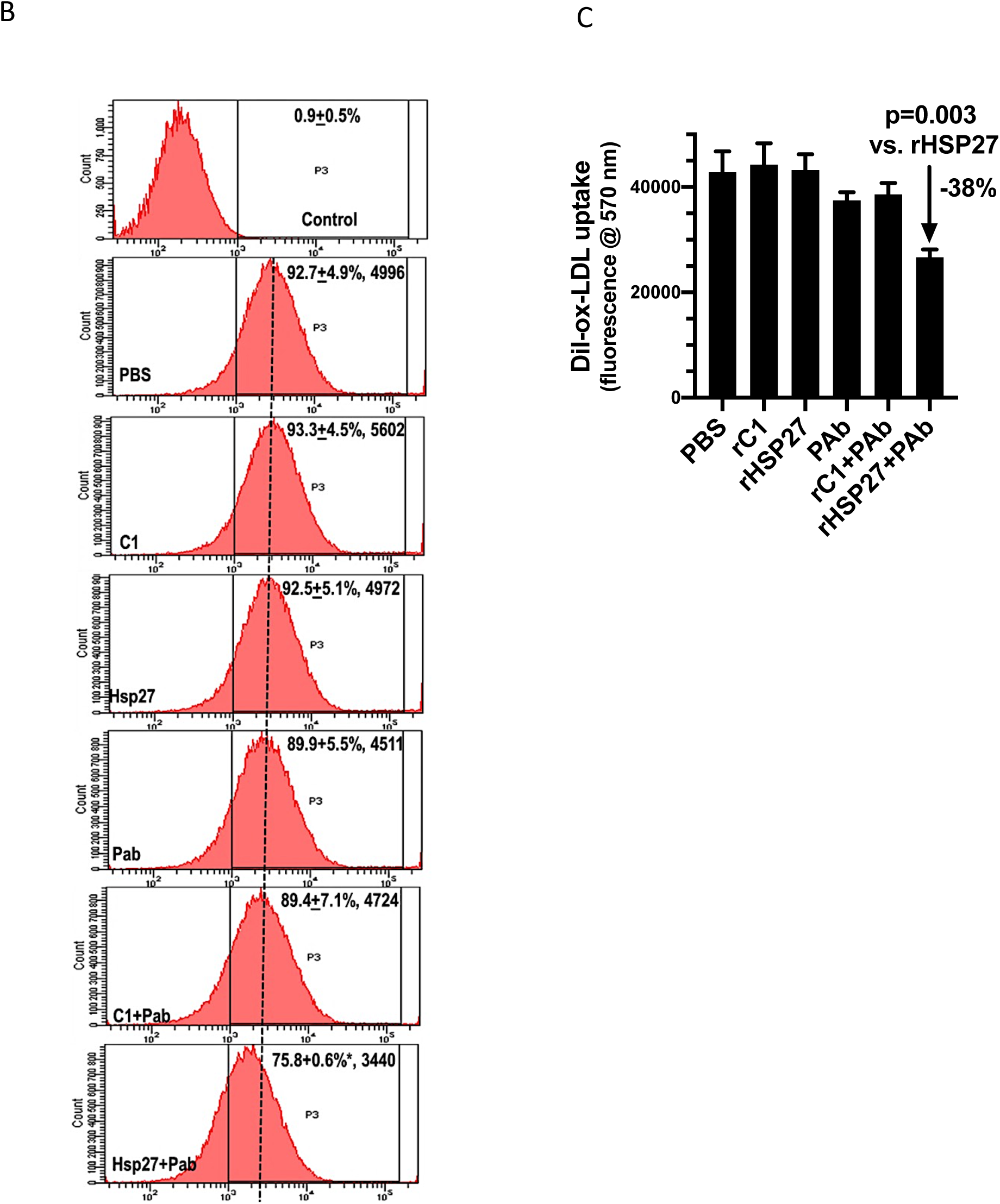
oxLDL uptake by scavenger receptors is inhibited by the HSP27 IC. A) Flow cytometry study of DiI-oxLDL uptake in HEK Blue™ cells expressing SR-AI (left), CD36 (middle) or Null (right). Cells were treated with combinations of rHSP27, rC1 and PAb, and the following parameters were recorded: i) percentage of cells positive for DiI-oxLDL fluorescence, and ii) The mean fluorescence intensity for DiI-oxLDL uptake per treatment group. Compared to PBS control, the presence of HSP27 IC reduced DiI-oxLDL uptake by 61% and 37% from SR-AI and CD36 expressing cells, respectively, whereas the other control treatments had negligible effects. B) THP-1 MΦ study similar to A), showing that compared to PBS control, Dil-oxLDL uptake is reduced from 92.7% to 75.8% with the HSP27 IC. Similarly, the MFI signal for the HSP27 IC was 31% lower than that of rHSP27 alone. C) Plate reader assay with THP-1 MΦ showing a 38% reduction in DiI-oxLDL uptake with the HSP27 IC compared to rHSP27 alone (p=0.003).
ii. Similarly, FACS was used to assess oxLDL uptake in THP-1 MΦ incubated with 1µg/ml rHSP27 or rC1 conjugated with FITC in the presence or absence of 5 µg/ml PAb (as well as PAb alone) for 16 hours. MΦ were then washed twice in PBS, detached using 1× trypsin-EDTA, washed in 1× PBS and the FITC signal was analyzed at 488 nm on a BD LSRII flow cytometer (**Fig. 3B**).
iii. Finally, using a plate reader assay, THP-1 MΦ (1 M/ml) grown to 80% confluence in 96 well plates and incubated for 2 hours with 1µg/ml DiI-oxLDL plus rHSP27 or rC1 with or without PAb, or the PAb alone. Plates were washed thrice with 200 µl DPBS, and the density of Dil-oxLDL was read using a BioTek Synergy Mx microplate reader at 570 nm (**Fig. 3C**).

### Assessment of rHSP27 Uptake by MΦ

Similar to the experiments described above, the uptake of FITC-labelled rHSP27 with or without PAb was assessed in MΦ using both FACs and phase contrast / fluorescent confocal microscopy (Fluoview FV10i Confocal Microscope; Olympus, Center Valley, PA) (**Fig. 4A – 4C**).

**Figure 4.**
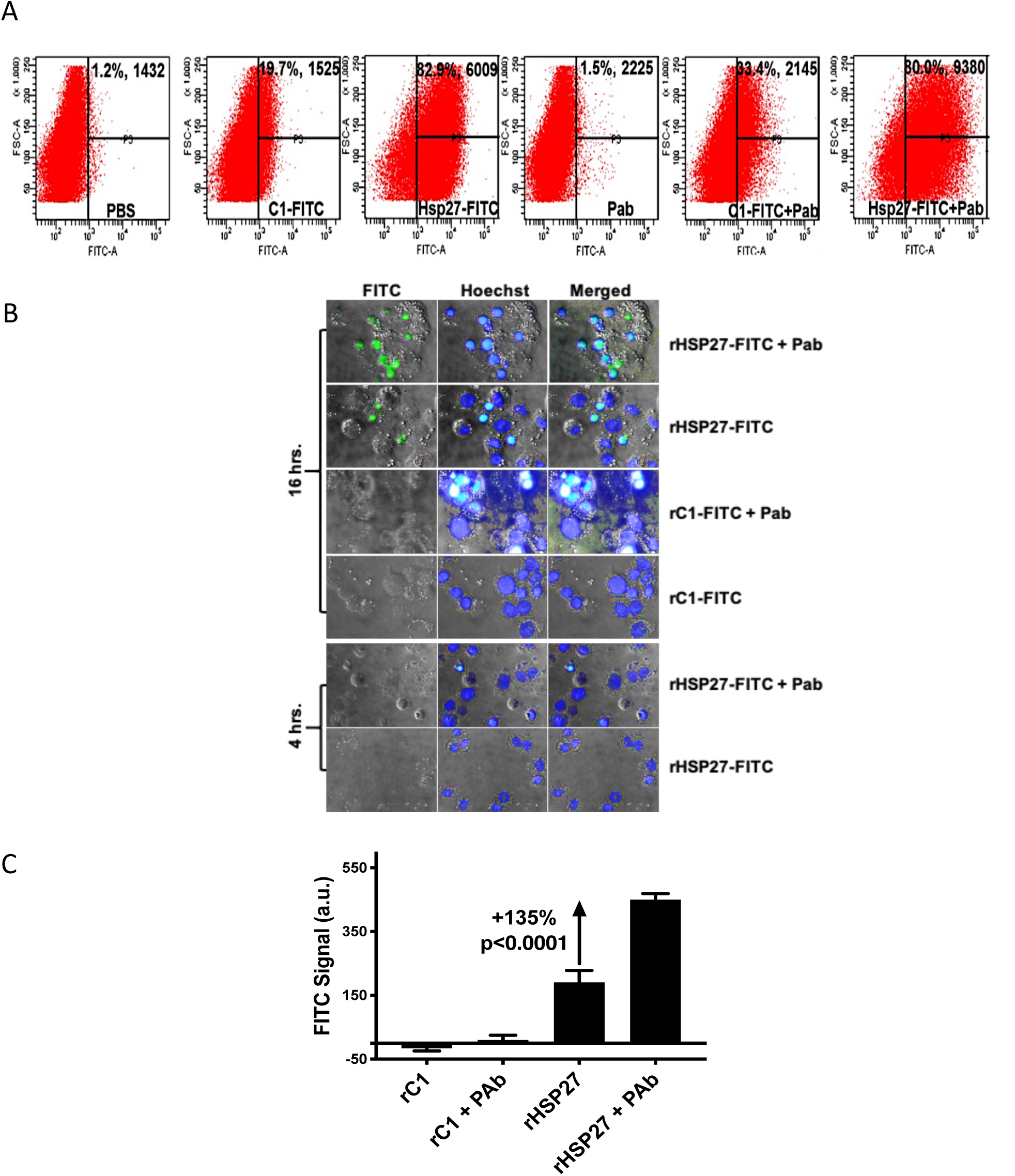
PAb enhances rHSP27 endocytosis in THP-1 MΦ. A) Using flow cytometry (A) the uptake of FITC-labelled rHSP27 is enhanced by with the addition of the PAb. B) Confocal microscopy demonstrating enhanced uptake of FITC-labelled rHSP27 in THP-1 MΦ at 16 but not 4 hrs (phase contrast and fluorescent microscopy; ×200). C) FITC-rHSP27 uptake is markedly increased when the PAb is combined with rHSP27.

### Statistical analysis

Unless stated otherwise, all data sets are represented as means ± standard error of the mean of experiments performed at least 3 times. Statistical significance was assessed by using a one-way ANOVA (Graph Pad Prism 8, La Jolla,CA, USA), and Tukey’s comparison of the groups. A p-value < 0.05 was considered significant.

## Results

### Characterization of the HSP27-AAb complex

A previously described rabbit polyclonal IgG anti-HSP27 antibody (PAb) with immunoreactivity similar to that of a commercial (goat) anti-HSP27 antibody and an epitope mapping pattern comparable to that of AAbs found in human serum was used for these studies (11). To characterize the size of the ICs formed with the addition of the PAb, rHSP27 and rC1 were incubated with PAb and the molecular sizes of the resulting ICs were determined by gel filtration analysis using FPLC (**Supplemental Fig. 1B**). The HSP27 IC is large (>30,000kDa) compared to rHSP27 alone (100-500kDa) or the PAb alone (∼200kDa). Increasing the rHSP27:PAb ratio from 1:1 to 1:5 led to a further increase in the IC size. The interaction between the PAb and rHSP27 was assessed using isothermal titration calorimetry and found to be an exothermic reaction (△H = -3.0 × 10^−4^ cal/mol) with a Ka of 8.9 ×10^−4^ M^-1^ with an average N valve of 0.5 – consistent with a 2:1 binding ratio for PAb:HSP27 (**Supplemental Fig. 1C**).

### Binding of the HSP27 Immune Complex to MΦ Cell Membrane

Recently, we showed that the HSP27 IC can alter gene expression (11).Hence, we now begin to explore its interaction with the cell membrane and signaling. While rHSP27 bound to the MΦ cell membrane with greater affinity than control treatments (BSA or rC1 without or with PAb), the PAb augmented rHSP27 cell surface binding 271% compared to rHSP27 alone (p<0.0001; **Fig. 1A**).

Given that exogenous HSP27 therapy attenuates the early inflammatory stages of atherogenesis (8, 10, 12), and that more recently this anti-inflammatory effect is amplified with rHSP25 vaccination (11), we sought explore the interaction between the HSP27 IC with TLR4. TLR4 protein was pulled down from the membrane fraction of THP-1 MΦ and verified using a monoclonal anti-TLR4 antibody confirmed to be specific to TLR4 (**Supplemental Fig. 2B**). Binding of biotinylated rHSP27 or rC1 (in the presence or absence of PAb) to TLR4 was assessed using strep-HRP activity to reflect the interaction intensity. The PAb promoted the interaction between rHSP27 and the TLR4 complex by 104% more than rHSP27 alone (**Fig. 1B**). rC1 alone or samples without either the anti-TLR4 antibody or the MΦ membrane protein fraction containing TLR4 showed only background noise signals. Interestingly, the interaction between [rC1 + PAb] and TLR4 was comparable to that of rHSP27 (alone) and TLR4 – perhaps indicating that it is the HSP27 alpha-crystallin domain (found in the rC1 fragment) that is important for the interaction with TLR4. The PAb did not interact with TLR4 complex, thus implying that the increased interaction with TLR4 in the presence of PAb was not from the PAb *per se*, but from the [rHSP27 + PAb] IC. Of note, the interaction with TLR4 increased by 125% as the ratio of HSP27 to PAb was increased from 1:1 to 1:5 (**Supplemental Fig. 2C**).

### PAb potentiates HSP27 NF-κB signaling in THP-1 MΦ

Previously, we show that rHSP27 activates NF-κB signaling in THP-1 MΦ and characterized the downstream transcriptional events. It is important to note that in these experiments we noted altered expression of both pro- and anti-inflammatory genes (12, 13). To determine if the binding of the HSP27 IC to TLR4 altered NF-κB signaling, we employed a SEAP reporter gene assay in HEK293 cells stably expressing TLR4 and its co-receptors, MD-2 and CD14 (i.e., HEK Blue™ TLR4 cells). HEK Blue™ parental cells deficient of TLR4 were used as a control (i.e., HEK Blue™ Null cells) and essentially showed no response to any treatment (**Fig. 1C**). In contrast to rHSP27 alone, the combination of [rHSP27 + PAb] showed 68% more NF-κB activation. LPS alone showed an NF-κB activation signal that was 122% greater than the HSP27 IC. Hence, TLR4 is required for HSP27 activation of the NF-κB pathway in these HEK Blue™ cells but, compared to LPS, was quite modest. It is important to note that both rC1 and rHSP27 are generated in bacteria – so if there was significant endotoxin contamination that might activate the NF-κB pathway it should be reflected by both of these treatments and therefore balance out. Moreover, we previously showed that adding polymyxin B to neutralize the effect of any potential endotoxin contaminants of the recombinant proteins made no difference in the NF-κB signal (12, 13).

Interestingly, when HEK Blue™ cells are treated with both LPS and [rHSP27 + PAb], NF-κB activation is moderately attenuated. For example, while treatment with LPS 10 and 100 ng/ml increased NF-κB activity by almost 4- and 6-fold; respectively. In the presence of the HSP27 IC the LPS-induced increase in NF-κB activity was reduced 26% and 46%; respectively (p<0.0001; **Fig. 1D**). The same concentrations of rC1, rHSP27 alone, PAb or [rC1 + PAb] showed neither inhibition nor promotion of NF-κB signaling compared to PBS control. Hence, these data suggest a competitive relationship between the HSP27 IC and LPS for the TLR4 receptor complex.

To further address this competition between LPS and the HSP27 IC, we set up a competitive binding assay for TLR4 pulled down from the membrane fraction of THP-1 MΦ. Compared to rHSP27 alone, [rHSP27 + PAb] reduced LPS binding to TLR4 by 77% (p<0.0001; **Fig. 1E)**. Finally, we assessed the downstream effects of these binding experiments by looking at THP-1 MΦ cytokine production. LPS-treated MΦ treated with [rHSP27 + PAb] released 167% higher levels of the anti-inflammatory cytokine IL-10 and 75% lower levels of the pro-inflammatory cytokine IL-1β compared to PBS control (p<0.001; **Fig. 1F**).

### The HSP27 Immune Complex interacts with MΦ Scavenger Receptors

Previously, we showed that rHSP27 (alone) binds SR-AI to reduce modified LDL uptake (9) and now assess if the addition of the PAb to rHSP27 alters oxLDL binding to SR-AI or its related scavenger receptor, CD-36. MΦ membrane extracts or SR-AI and CD-36 were pulled down and their identities were verified by Western blotting (**Supplemental Fig. 2B)**. Compared to rHSP27 alone the PAb promoted the interaction between rHSP27 and SR-AI, as well as rHSP27 and CD-36 (p<0.01 and p<0.001 respectively; **Fig. 2A, 2B**). rC1 alone, PAb alone or samples without either the anti-SR-AI or anti-CD-36 antibodies, or the MΦ membrane protein fraction containing either SR-AI or CD-36 showed only background noise signals. Interestingly, the interaction between [rC1 + PAb] and CD-36 was comparable to that of rHSP27 (alone) – perhaps indicating that it is the HSP27 alpha-crystallin domain (also found in the rC1 fragment) that is important for the interaction with CD-36. Of note, as the ratio of HSP27 to anti-HSP27 antibody is increased from 1:1 to 1:5, using excess PAb, the interaction with SR-AI and CD-36 increased by approximately 2.5- and 1.5-fold; respectively (**Supplemental Fig. 2D, 2E**).

### HSP27 Immune complex competes with oxLDL for SR-AI and CD36 binding

As both scavenger receptors are involved in MΦ lipoprotein uptake we next assessed whether the HSP27 IC can compete for oxLDL binding to SR-AI or CD-36 pulled down from MΦ cell lysates. [rHSP27 + PAb] was superior to rHSP27 alone in competing with oxLDL for binding to SR-AI or CD36, with less than 0.5 μg/ml rHSP27 combined with PAb almost completely nullifying oxLDL binding to either receptor. (**Fig. 2C, 2D**). The converse experiment was also carried out, using increasing concentrations of oxLDL to try to displace rHSP27 binding to either scavenger receptor. Essentially, rHSP27 combined with PAb showed ≥95%% binding to SR-AI or CD-36 regardless how high the concentration oxLDL was increased (to a maximum of 164 μg/ml; **Fig. 2E, 2F**)). In contrast, rHSP27 alone was displaced from SR-AI and CD-36 at the maximum oxLDL concentration (e.g., 28% and 14% reductions in rHSP27 binding; respectively). Together, these results show that the PAb potentiated the interaction of rHSP27 with both SR-AI and CD36, thereby attenuating oxLDL binding to its cognate receptors.

### HSP27-PAb complex attenuate uptake of oxLDL

To assess the relative contribution of SR-AI and CD36 in oxLDL uptake we used HEK Blue™ Null1-v cell line which is devoid of both of these receptors, and then separately transfected SR-AI and CD-36 into these parental cells. First, the expression of these scavenger receptors and their abundance in HEK Blue™ cells relative to THP-1 MΦ was confirmed using both Western blotting and FACS (**Supplemental Fig. 3A – 3D**). HEK Blue™ cells expressing SR-AI or CD-36 were then treated with DiI-oxLDL and combinations of rHSP27, rC1 and PAb for 2 hrs before performing FACS. For each experimental condition, the percentage of fluorescent cells was measured, as well as the MFI, a reflection of the amount of oxLDL that was taken up per experimental condition. HEK Blue™ Null-v cells served as negative controls and showed low (background) levels of fluorescence for PBS and all other treatments, thereby highlighting the requirement for SR-AI and CD-36 for oxLDL uptake.

For HEK Blue™ SR-AI cells the percentage of oxLDL positive cells was similar for all control conditions (PBS: 97.1%, rC1: 97.7%, PAb: 97.4%), as well as treatment with rHSP27 alone (97.5%). In contrast, treatment with the combination of [rHSP27 + PAb] resulted in 81.0% of the cells being oxLDL positive, reflecting a modest decrease. While the MFI was similar amongst the control treatments, oxLDL uptake with HSP27 IC treatment was 65.2% lower than with rHSP27 treatment (2,390 *vs*. 6,869 a.u.) (**Fig. 3A**).

For the HEK Blue™ CD-36 cells, the percentage of cells with oxLDL uptake for the control treatments were similar (e.g., PBS 92.2%, rC1 91.8%, PAb 92.9%). However, the percentage of cells with oxLDL uptake for the [rHSP27 + PAb] treatment group was only 73.3% compared to the cells treated with rHSP27 alone (93.0%). For the cells treated with [rHSP27 + PAb] the MFI was 43.3% lower than those treated with rHSP27 alone (4,815 *vs*. 2,731 a.u., **Fig. 3A**).

Next we addressed the same question of the effect of the HSP27 IC on oxLDL uptake in THP-1 MΦ using both FACS and a plate reader assay. DiI-oxLDL uptake was reduced in the presence of HSP27 IC by both techniques:

a. For the FACS studies, the percentage of cells fluorescent for DiI-oxLDL was similar amongst the controls: PBS (92.7%), rC1 (93.3%) and PAb (89.95), but dropped to 75.8% with [rHSP27+PAb] treatment (**Fig. 3B**). With regards to the MFI signal that reflects the amount of DiI-oxLDL uptake for each experimental condition, the rHSP27 (alone) treatment group (4,972) was indistinguishable from the PBS control (4,996 a.u.). However, MΦ treated with [rHSP27 + PAb] had a 31% lower MFI signal (3,440 a.u) compared to rHSP27 alone.
b. A plate reader assay showed a similar 38% reduction of DiI-oxLDL uptake with the rHSP27 IC compared to rHSP27 alone (p=0.003), and no changes with rC1, rHSP27, PAb alone or rC1 and PAb (**Fig. 3C**).

### rHSP27 Internalization

FACS studies were also used to examine the uptake of rHSP27 when the PAb is added. The percentage of cells that were fluorescent for FITC-labeled rHSP27 was comparable amongst the controls: PBS (1.2%) and PAb (1.5%); while, the uptake of rC1 alone (19.7%) and rC1+PAb (33.4%) were higher. Nonetheless, rHSP27 alone (82.9%) and [rHSP27+PAb] (80.0%) were much higher, and similar – thereby suggesting that approximately the same percentage of cells showed uptake of rHSP27 *vs*. the rHSP27 IC. However, on reviewing the amount of FITC-labelled rHSP27 taken up (i.e., the MFI), treatment with the rHSP27 IC resulted in 56.1% greater uptake relative to rHSP27 alone (9,380 vs. 6,009), with all controls being much lower (**Fig. 4A**). A rHSP27 uptake assay using a plate reader to quantify the FITC signal in MΦ showed virtually no uptake at 4 hrs, but by 16 hrs the uptake was 135% greater with the HSP27 IC compared too rHSP27 alone (p<0.001), with negligible uptake of rC1 with or without of the PAb (**Fig. 4B, 4C**). Like the FACS result, there did not seem to be an overwhelming increase in the number of FITC positive MΦ, rather the brightness signal in the positive cells appeared stronger, thereby implying increased uptake in certain but not all cells.

## Discussion

Investigations by several laboratories, including our own, highlight the point that HSP27 arterial expression and blood levels are higher in health compared to cardiovascular disease (3-7). As well, we now know that AAb levels are also higher in health but have yet to fully understand their role (11). Recently, we generated data to suggest that [HSP27 + PAb] can act from outside the cell to produce signals that result in reduced inflammation and blood cholesterol levels due to increased expression of the low density lipoprotein (LDL) receptor that requires TLR4 and activation of the NF-κB pathway (11). A second major series of observation from our group is that interventions that increase HSP27 levels are associated with reduced plaque content of both MΦ and cholesterol clefts (8, 10). We postulate that this effect is not solely due to reduced plasma cholesterol levels, as previously we showed that HSP27 (alone) can bind SR-AI to reduce uptake of modified LDL particles *in vitro* (19), we have not taken into account the potential role of AAbs. Similarly, a number of important studies from eminent groups in the heat shock protein (HSP) field provide important insights into how various HSPs, including HSP27, interact at the cell membrane (20-23), yet none consider the potential role of antibodies to the respective HSPs. Intuitively, one might surmise that these anti-HSP27 antibodies are detrimental to the biology of HSP27, essentially covering up the protein and barring it for performing its beneficial extracellular functions. Hence, the goal of this study was to examine the effect of combining a validated anti-HSP27 polyclonal IgG antibody with rHSP27 on the TLR4/NF-κB pathway, as well MΦ scavenger receptor biology. Three important clusters of observations arose from these studies and now form the basis of a revised working model: HSP27 **<u>I</u>**mmune **<u>C</u>**omplex **<u>A</u>**ltered **<u>S</u>**ignaling and **<u>T</u>**ransport (ICAST; **Fig. 5**).

**Fig. 5.**
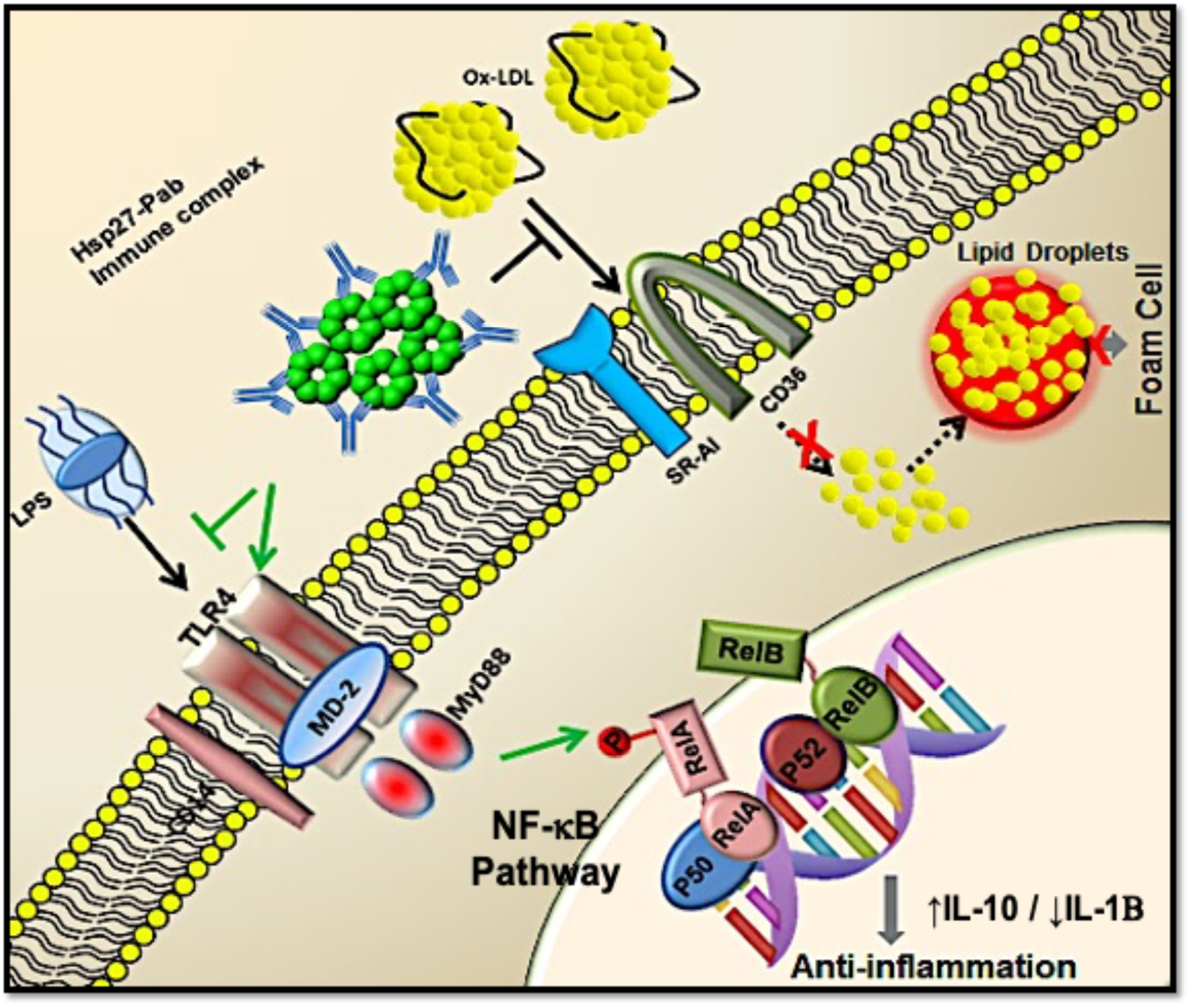
ICAST Schematic. Outline of the working hypothesis as to how the HSP27 IC reduces MΦ inflammatory signaling, blocks oxLDL uptake and can undergo internalization. Binding of HSP27 to the cell membrane is facilitated by the addition of the AAb. The HSP27 IC interacts with TLR4 to activate the NF-κB pathway, competitively displacing the interaction of LPS with this receptor. Activation of NF-κB by this external pathway effects an anti-inflammatory milieu with levels of secreted IL-10 increased and IL-1β decreased. As well, the HSP27 IC competes with oxLDL for binding to scavenger receptors SR-AI and CD-36, thereby reducing oxLDL internalization and foam cell formation. Finally, the presence of the PAb enhances the internalization of HSP27 (not shown) via mechanisms that are currently under study.

First, the HSP27 IC is superior to HSP27 alone in binding the MΦ membrane (**Fig. 1**). TLR4 is important in the interaction of the HSP27 IC with the MΦ membrane, and is definitely required for the activation of NF-κB. The transcription factor NF-κB is a key regulator of inflammation, immune responses, cell survival, and cell proliferation (24). Upon activation, NF-κB can mediate the induction of more than 160 genes, many of which have documented roles in both atherogenesis and athero-protection (12, 25). Activated NF-κB has been detected in endothelial cells, smooth muscle cells, and MΦ in atherosclerotic plaques, which may be involved in either the development or protection of atherosclerosis (26, 27). Interestingly, we note in this study a decrease in LPS-mediated NF-κB activation when the HSP27 IC is present. Moreover, treatment of MΦ with the HSP27 IC markedly reverses the balance of the inflammatory cytokines secreted in response to LPS treatment (e.g., a 75% reduction in IL-1β and a 167% increase in IL-10). So there appears to be a yin and yang, whereby activation of NF-κB can be detrimental if it results in primarily pro-inflammatory effects (e.g., LPS alone). Alternatively, triggering the NF-κB pathway with HSP27 ICs can suppress (or out-compete) inflammatory mediators such as LPS for TLR4, and/or enact athero-protective effects, such as the upregulation of LDLR transcription (11). Certainly, this dualism in HSP27 IC signaling is becoming increasing interesting and forms the basis of our future studies.

Second, the HSP27 IC binds both SR-AI and CD-36, and can displace oxLDL binding with these scavenger receptors, resulting in less oxLDL uptake in MΦ (**Fig. 2, 3**). Out of all of the handful of scavenger receptors that have been identified, SR-AI and CD36 are the two main receptors interacting with and then ingesting oxLDL.(28) oxLDL plays a key pro-atherogenic role in the arterial wall. Various known vascular risk factors are associated with increased oxidative stress that transforms native LDL to oxLDL and the formation of foam cells that populate atherosclerotic plaques. Previously we showed that the uptake of oxLDL by THP-1 MΦ is hampered by treatment with high dose HSP27 alone (250 µg/ml) (9, 13), which actually down-regulates SR-AI expression (13). In the current studies we note that at more physiological HSP27 concentrations (1 µg/ml) there is no inhibition of MΦ oxLDL uptake, except when rHSP27 combines with the PAb. Indeed, compared to HSP27 alone, the HSP27 IC has a stronger affinity for both SR-AI and CD36, thereby out-competing oxLDL and reducing its endocytosis through these receptors.

Third, combining the PAb with rHSP27 results in enhanced cellular uptake and internalization of rHSP27 in MΦ. Although we recently noted how HSP25 vaccination in mice augments the expression of LDLR by increasing the activity of its promoter (11), we struggled with trying to understand how HSP27 could somehow traverse the cell membrane to alter transcription. The current data provide initial insights regarding the internalization process, highlighting how it is facilitated by the presence of the anti-HSP27 antibody. Certainly, more study is required to dissect the internal signaling pathways before we can better understand the effects of HSP27 on (say) LDLR transcription, and if there are direct or indirect effects on the LDLR promoter.

In summary, the IC formed between HSP27 and its autoantibody, potentiates the interaction with the MΦ cell membrane, a process that involves TLR4 and the scavenger receptors SR-AI and CD-36. The activation of the NF-κB pathway is more potent with the HSP27 IC *vs*. HSP27 alone, yet modest compared to LPS alone. Similarly, the HSP27 IC is superior to HSP27 alone in competing with SR-AI and CD-36 for the uptake of oxLDL – which may be very important for not only plaque MΦ, but also the in the development of non-alcoholic fatty liver disease, an important metabolic issue that predisposes to liver fibrosis and coronary artery disease. To date, this is the first report of the HSP27 IC acting as an innate antagonist of inflammatory signaling and MΦ foam cell formation. Taken together, these studies provide further stimulus to develop either active or passive HSP27 immunization strategies that boost AAb levels for the prevention and/or treatment of atherosclerosis and other inflammatory diseases.

## Nonstandard Abbreviations

AAb: anti-HSP27 (or anti-HSP25) antibodies
ApoE^-/-^: apolipoprotein E null
a.u.: absorbance units
BSA: bovine serum albumin
HSP: heat shock protein
HSP27: Heat Shock Protein 27
IL-10: interleukin-10
IL-1β: interleukin-1 beta
LDL: low density lipoprotein cholesterol
LDLR: low density lipoprotein receptor
NF-κB: nuclear factor kappa light chain enhancer of activated B cells
ON: overnight
oxLDL: oxidized LDL
PBS: phosphate buffered saline
PBST: PBS tween
rC1: recombinant C-terminal truncation of HSP27 spanning amino acids 90-205
rHSP27: recombinant HSP27
Strep-HRT: streptavidin-horse radish peroxidase
TLR-4: toll-like receptor 4
TMB: 3,3′,5,5′-tetramethylbenzidine

## Acknowledgments

We are indebted to the Libin Cardiovascular Institute of Alberta and its community partners for supporting the O’Brien Vascular Biology Laboratory and associated infrastructural research entities. We appreciate the support of many collaborators at the University of Calgary including: Paul Kubes PhD for TLR4^-/-^ mice, Dr. Elmar Prenner’s laboratory in the Department of Biological Sciences where the ITC studies were performed and Yiping Liu, MD, PhD in the Flow Cytometry core facility at the Cumming School of Medicine.

## Notes

### Competing Interest Statement

EOB is the inventor on US Provisional Patent Application (serial number: 63/031,640). EOB is the Scientific Co-Founder of Pemi31 Therapeutics Inc., a startup company that controls the aforementioned intellectual property. EOB, CS and YXC have equity interests in Pemi31 Therapeutics Inc.

