## Supplemental Figures & Legends for "Heat Shock Protein 27 Immune Complex Altered Signaling and Transport (ICAST): Novel Mechanisms of Attenuating Inflammation"

Supplemental Fig. 1

A

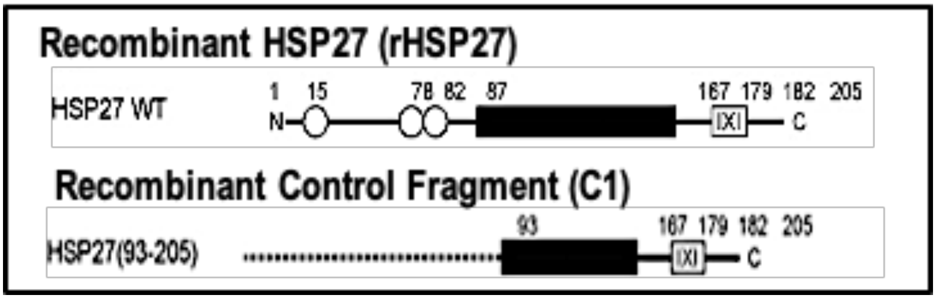


B

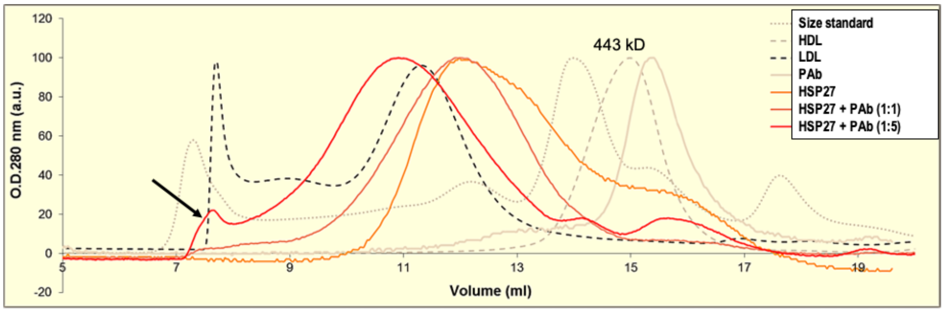


C

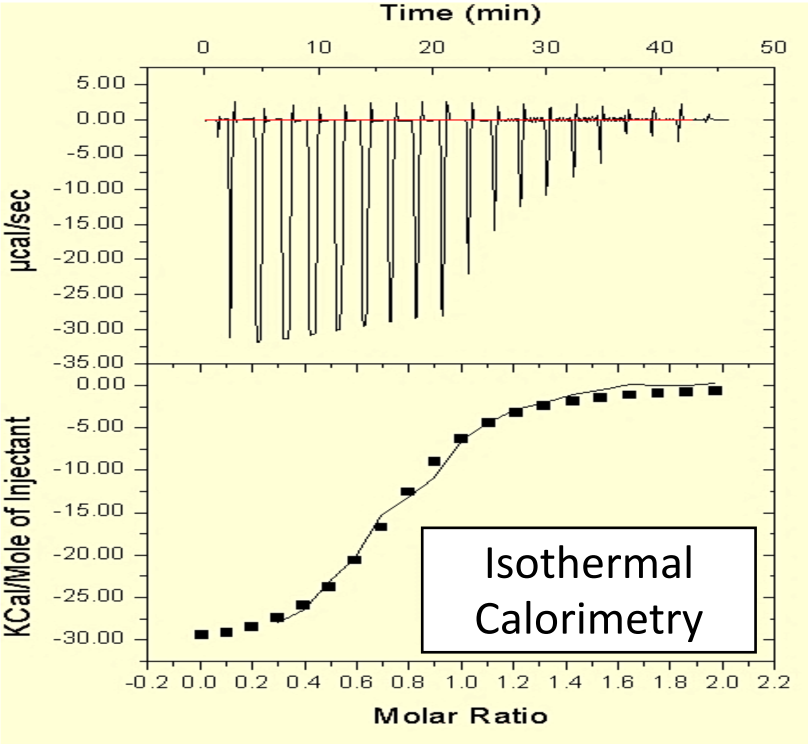

**Supplementary Fig. 1. Characterization of the HSP27 IC**

A) Schematic showing the full-length protein sequence of rHSP27 and the truncated rC1 fragment that spans a region of the protein containing the β-crystallin domain in the C-terminus.

B) Size exclusion chromatography showing the much larger size of the HSP27 IC compared to HSP27 alone and other moieties. Increasing the rHSP27:PAb ratio from 1:1 to 1:5 led to a further increase in the IC size (arrow). The size standard has a dominant peak at 443 kD, representing apoferritin.

C) Isothermal titration calorimetry demonstrating a strong, exothermic interaction between the PAb and rHSP27.

Supplemental Fig. 2

A

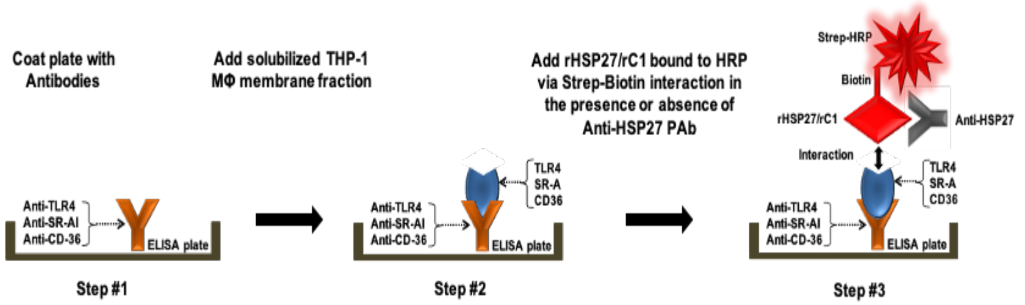


B

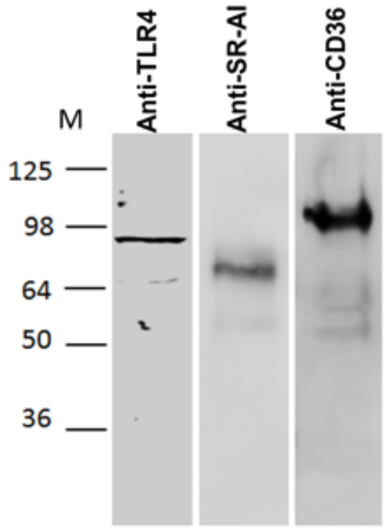


C

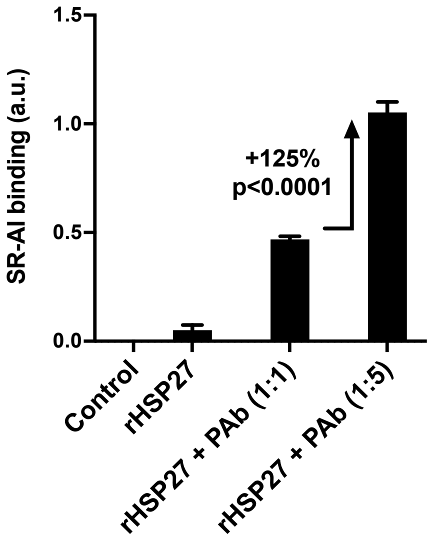


D

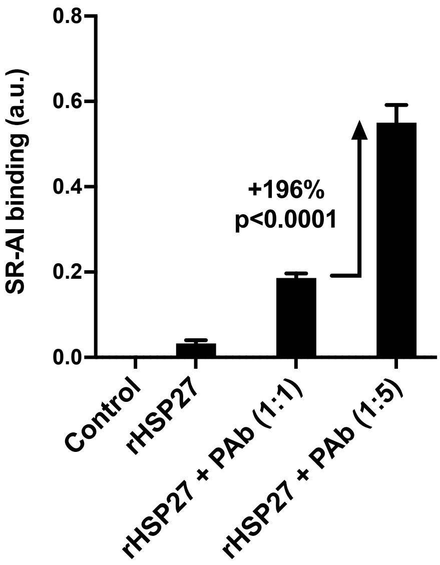


E

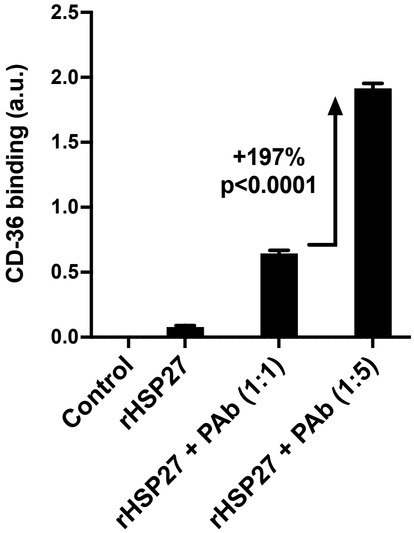

**Supplementary Fig. 2.** ***In vitro*** **binding assay for TLR4, SR-AI and CD-36 with HSP27 IC**

A) Schematic diagram showing the steps involved in the binding assays for TLR4, SR-AI and CD-36.

B) Western blot showing specificity of monoclonal antibodies to TLR4, SR-AI, or CD36 that were used for the *in vitro* binding assays.

C) – E) The higher HSP27:PAb ratio (1:5) improved the interaction of rHSP27 with TLR4 (C), SR-AI (D), and CD36 (E) compared to a 1:1 rHSP27:PAb ratio.

Supplemental Fig. 3

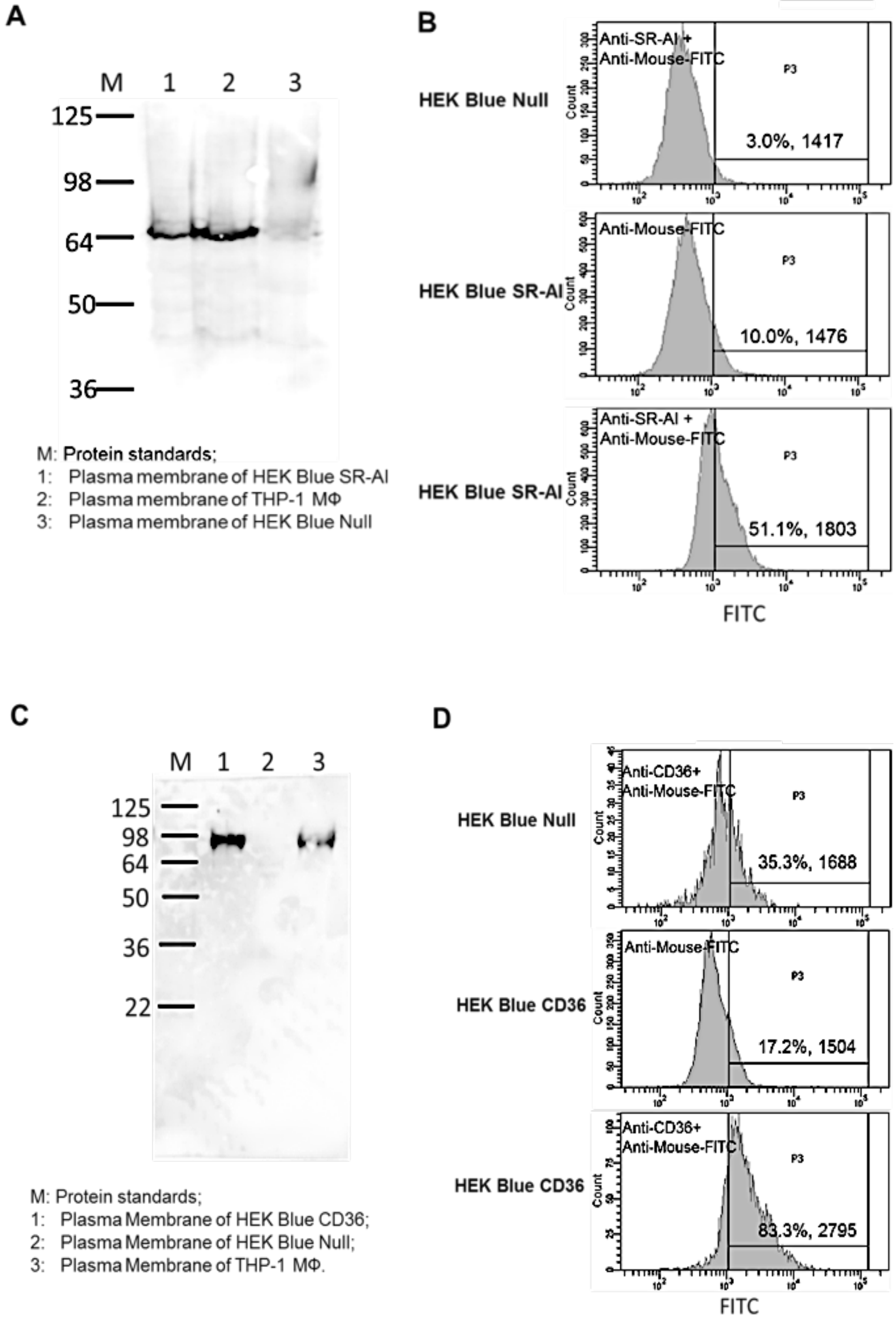

**Supplementary Fig. 3. Validation of SR-AI and CD36 Expression in HEK Blue Null-1 V cells**

A) Western blot of plasma membrane fractions of HEK Blue Null cells expressing SR-AI (Lane 1), plasma membrane fraction of THP-1 MΦ as the positive control (Lane 2) and plasma membrane of HEK Blue Null cells as the negative control (Lane 3).

B) Flow cytometry of HEK blue cells to confirm the expression of transfected SR-AI. Top panel: HEK Blue Null cells served as a negative control. Middle panel: HEK Blue SR-AI cells immunolabeled with an anti-mouse FITC antibody but no anti-SR-AI antibody (i.e., another negative control). Bottom panel: using anti-SR-AI as the primary antibody and anti-mouse IgG-FITC as the signal antibody to demonstrate the presence of SR-AI in 51% of cells.

C) Western blot of plasma membrane fractions of HEK Blue Null cells expressing CD36 (Lane 1), plasma membrane fraction of THP-1 MΦ as the positive control (Lane 3) and plasma membrane of HEK Blue Null cells as the negative control (Lane 2).

D) Flow cytometry of HEK blue cells to confirm the expression of transfected CD-36. Top panel: HEK Blue Null cells served as a negative control. Middle panel: HEK Blue CD36 cells immunolabeled with anti-mouse FITC antibody but no anti-CD36 antibody (i.e., another negative control). Bottom panel: using anti-CD36 as the primary antibody and anti-mouse IgG-FITC as the signal antibody to demonstrate the presence of CD36 in 83% of the cells.
